# The DNMT1 inhibitor GSK-3484862 mediates global demethylation in murine embryonic stem cells

**DOI:** 10.1101/2021.09.12.459949

**Authors:** Nathalia Azevedo Portilho, Deepak Saini, Ishtiaque Hossain, Jacinthe Sirois, Christopher Moraes, William A. Pastor

## Abstract

**Background:** DNA methylation plays an important role in regulating gene expression in mammals. The covalent DNMT1 inhibitors 5-azacytidine and decitabine are widely used in research to reduce DNA methylation levels, but they impart severe cytotoxicity which limits their demethylation capability and confounds interpretation of experiments. Recently, a non-covalent inhibitor of DNMT1 called GSK-3484862 was developed by GlaxoSmithKline. We sought to determine whether GSK-3484862 can induce demethylation more effectively than 5-azanucleosides. Murine embryonic stem cells (mESC) are an ideal cell type in which to conduct such experiments, as they have a high degree of DNA methylation but tolerate dramatic methylation loss.

**Results:** We determined the cytotoxicity and optimal concentration of GSK-3484862 by treating wild-type (WT) or *Dnmt1/3a/3b* triple knockout (TKO) mESC with different concentrations of the compound, which was obtained from two commercial sources. Concentrations of 10 µM or below were readily tolerated for 14 days of culture. Known DNA methylation targets such as germline genes and GLN-family transposons were upregulated within two days of the start of GSK-3484862 treatment. By contrast, 5-azacytidine and decitabine induced weaker upregulation of methylated genes and extensive cell death. Whole genome bisulfite sequencing (WGBS) showed that treatment with GSK-3484862 induced dramatic DNA methylation loss, with global CpG methylation levels falling from near 70% in WT mESC to less than 18% after 6 days of treatment with GSK-3484862, similar to the methylation level observed in *Dnmt1* deficient mESCs.

**Conclusions:** GSK-3484862 mediates striking demethylation in mESCs with minimal non-specific toxicity.

## Introduction

DNA methylation is a key regulator of gene expression in mammals[1]. DNA methyltransferases (DNMTs) transfer a methyl group to the fifth carbon of the DNA base cytosine to form 5-methylcytosine. Typically, a high density of DNA methylation at a gene or transposon’s transcriptional start site promotes transcriptional repression[2]. DNMT3A and DNMT3B methylate previously unmodified CpG sites, establishing the pattern of DNA methylation during early embryonic development[3]. DNMT1 in turn methylates the newly synthesized DNA strand after DNA replication. Because DNMT1 preferentially methylates CpGs complementary to, or in the vicinity of, existing CpG methylation, it can maintain CpG methylation patterns through many cycles of cell division[4, 5].

DNA methylation is critical for maintaining silencing of certain genes, particularly genes associated with germline development, and transposons[1, 6, 7]. Aberrant hypermethylation of the promoters of tumour suppressors and other genes occurs in a wide variety of cancers[8]. Consequently, there is high interest in demethylating drugs for both research and therapeutic purposes[9]. Two such drugs are widely used in research and as therapeutics: 5-azacytidine and decitabine (5-aza-2’-deoxycytidine). They were synthesized in the 1960s as potential chemotherapeutics that might interfere with nucleic acid metabolism in rapidly dividing cells [10, 11]. It was subsequently demonstrated that unlike other chemotherapeutics, 5-azacytidine and related compounds induce differentiation of sarcoma cells to myotubes, an activity attributable to apparent demethylating properties [12].

5-azacytidine and decitabine are cytidine analogs which contain nitrogen at the fifth position of the pyrimidine ring [10, 11]. In cells, decitabine is converted to 5-aza-dCTP and incorporated into DNA during replication[13]. The 5-azacytosine base forms a stable covalent bond with DNMT1, irreversibly inhibiting the enzyme[14]. The resulting DNA-protein adduct is then repaired and DNMT1 is destroyed in the proteosome[15]. Decitabine thus has two modes of activity: it causes cell death via the creation of excessive DNA-protein adducts, and it reduces DNA methylation by the inhibition and destruction of DNMT1[16]. 5-azacytidine is converted into a mixture of 5-aza-dCTP and 5-aza-CTP in the cell[17]. As such, it is incorporated into both DNA and RNA, and may cause additional cytotoxicity by interfering with RNA metabolism, methylation, and translation[18, 19].

The cytotoxicity of 5-azanucleosides limits their use as demethylating agents in research. Excessive concentrations of 5-azanucleosides will cause cell death, and thus experiments must be conducted in a narrow window of optimal concentration in which DNMT1 is inhibited but the cells do not die[9]. At tolerated doses, DNMT1 inhibition may be incomplete, and 5-azanucleoside-treated cells show only modest global reductions in DNA methylation when measured by highly quantitative whole genome bisulfite sequencing (WGBS)[20, 21]. Even at lower concentrations, the genotoxic effects of 5-azanucleoside make it unclear whether gene expression changes and cellular phenotypes are caused by demethylation or non-specific toxicity. To identify genes regulated by DNA methylation, Hackett and colleagues treated NIH3T3 cells with decitabine for 3 days and identified 344 upregulated genes[22]. Few of these upregulated genes had heavily methylated promoters and many upregulated genes were involved in immune and stress response, suggesting a response to toxicity rather than demethylation. After 14 days of recovery, 49 genes showed persistent upregulation. These genes were disproportionately enriched for heavily methylated promoters and likely to be direct targets of methylation-mediated silencing. This strategy of treatment and recovery was successful at identifying genes regulated by DNA methylation in NIH3T3s but carries obvious drawbacks. Any gene remethylated during the recovery period would be missed. Furthermore, researchers studying the role of DNA methylation during a dynamic process cannot necessarily incorporate a treatment and recovery period. A DNA methyltransferase inhibitor lacking non-specific toxicity would be preferable from a research perspective.

Despite their limitations, 5-azanucleosides have remained the best and most widely used drugs for inducing DNA demethylation for the last 40 years. In 2019, GlaxoSmithKline announced the discovery of a non-covalent DNMT1 inhibitor, GSK-3484862 [23]. As of this writing, there is no publication describing its initial discovery in detail, but it has been used in two articles. Researchers at GlaxoSmithKline demonstrate that GSK-3842364, a racemic mixture including GSK-3484862 and its enantiomer, selectively inhibits DNMT1 with an IC_50_ of 0.4 µM[24]. Five days of treatment of GSK-3842364 induces a dramatic global demethylation of erythroid progenitor cells as measured by mass spectrometry. In addition, GSK-3842364 shows lower cytotoxicity than decitabine and induces transcription of the methylated fetal hemoglobin genes, *HBG1* and *HBG2*, both in erythroid cells *in vitro* and in a murine model of sickle cell anemia. In the second report, Haggarty and colleagues used GSK-3484862 to inhibit DNMT1 in murine pre-implantation *Dnmt3a/3b* KO embryos[25]. Here, a concentration above 0.35 µM prevents blastocyst formation. At this low concentration, only a 34% global drop in DNA methylation is observed relative to untreated *Dnmt3a/3b* KO embryos; however, the treatment is sufficient to induce a marked increase in IAP-Ez transposon expression, consistent with demethylating activity of the compound.

Considering the limited data concerning GSK-3484862 and the importance of a better DNA methylation inhibitor for the scientific community, we tested the demethylating capacity of GSK-3484862 in murine embryonic stem cells (mESCs), which are ideal for testing methyltransferase inhibitors. When cultured in classic serum+LIF conditions, mESCs have a high global level of DNA methylation[26]. However, while most mammalian cells require some DNA methylation for survival[27], *Dnmt1/3a/3b* triple knockout (TKO) embryonic stem cells self-renew and proliferate normally[28]. The set of genes regulated by DNA methylation in mESCs is well characterized, as are individual and compound DNA methyltransferase knockouts[29, 30]. In addition to wild-type cells, we used *Dnmt1/3a/3b* TKO mESCs because lacking DNA methylation, these cells are suitable to assess non-specific cytotoxicity of GSK-3484862. Our results indicate that GSK-3484862 mediates striking demethylation in mESCs, comparable to what is observed in a complete DNMT1 knockout, with minimal non-specific toxicity.

## Results

### mESCs tolerate high concentrations of GSK-3484862

As of this writing, GSK-3484862 is not available for sale from GlaxoSmithKline, so we conducted experiments using preparations purchased from ChemieTek and MedChemExpress. First, we sought to determine the cytotoxicity and optimal concentration of GSK-3484862 (Fig. 1). Wild-type J1 line and *Dnmt1/3a/3b* triple knockout (TKO) mESCs on a J1 background were seeded at low density on gelatin-coated plates. WT mESCs showed classic mESC morphology, dome-shaped colonies with rounded edges, while TKO ESCs showed jagged colony edges when cultured on gelatin. Cells were treated 24 hours after plating with concentrations of GSK-3484862 ranging from 2 pM to 200 µM. Substantial precipitation of the drug in media was observed at concentrations equal to or greater than 20 µM. After six days the morphology of WT and TKO cells treated with GSK-3484862 was similar to that of corresponding control cells that received only DMSO, with mortality only noticeable at 200 µM (Fig.1A). No differences were observed in cell numbers after six days of treatment with GSK-3484862 at concentrations equal to or below 20 µM in WT and TKO cells. (Fig. 1B).

**Figure 1.**
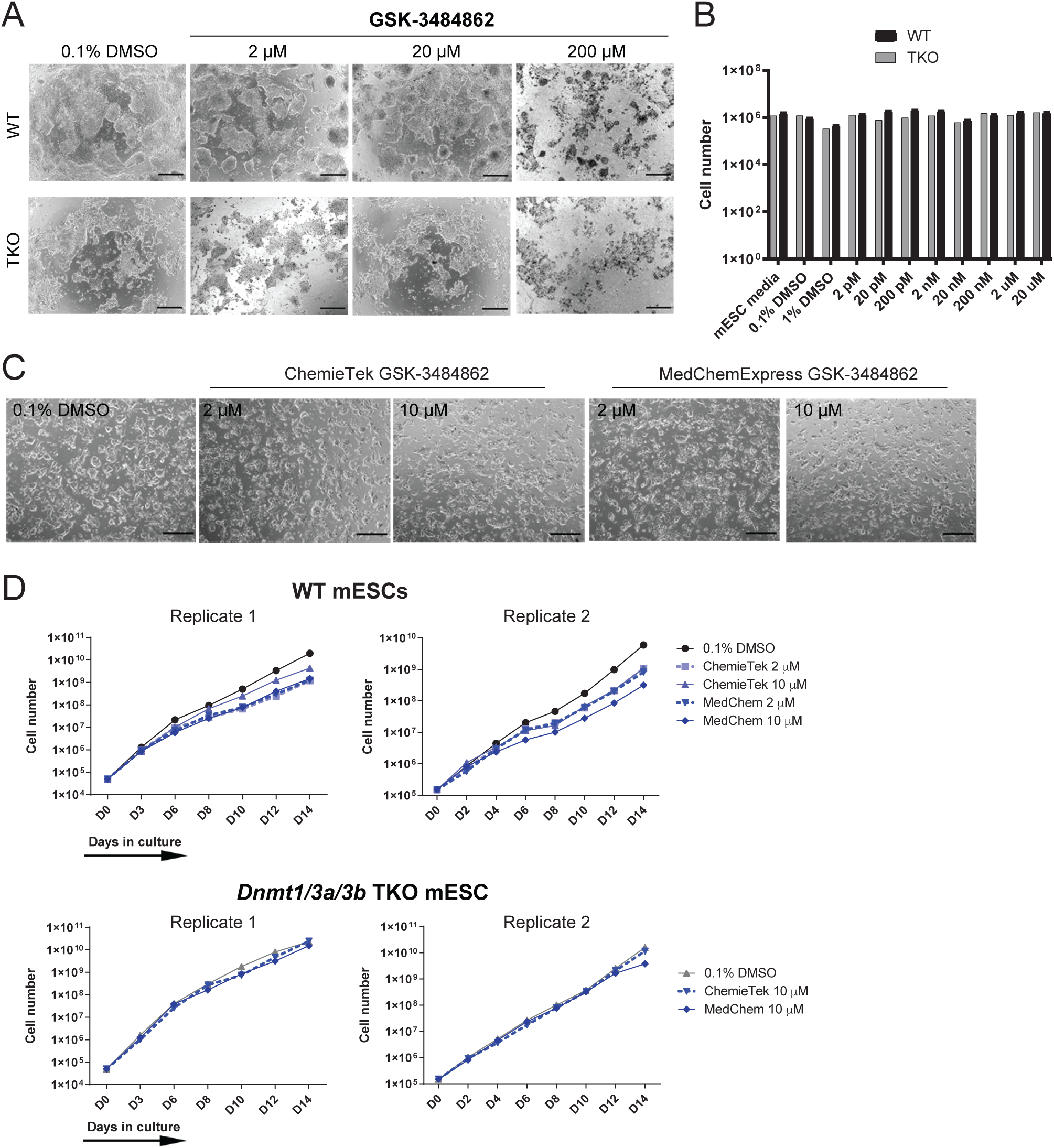
GSK-3484862 inhibitor treatment in mouse embryonic stem cells (mESC). **A** Brightfield images showing morphology of WT and TKO cells after 6 days of treatment with 0.1% DMSO or GSK-3484862 (2 µM, 20 µM and 200 µM) obtained from ChemiTek. **B** Cell numbers after 30,000 WT or TKO mESC were treated with indicated culture conditions for six days. **C** Brightfield images of WT cells after 14 daysof treatment with 0.1% DMSO, 2 µM or 10 µM of GSK-3484862 from ChemieTek or MedChemExpress. **D** Graphs from two independent experiments show the cell growth of WT or TKO cells counted every 2-3 days.

We then evaluated the long-term cytotoxicity and demethylation efficacy of GSK-3484862. WT and TKO cells were treated with 2 or 10 µM of GSK-3484862 for 14 days, 10 µM being the highest concentration at which drug precipitation was not observed. WT drug-treated mESCs were indistinguishable from corresponding vehicle-treated controls throughout the 14 days (Fig. 1C). A slight but reproducible decrease in cell growth was observed in GSK-3484862-treated WT cells from day 6 onwards, whereas the growth rate of TKO cells was constant throughout treatment (Fig. 1D). No difference was observed between compound produced by the two companies (Fig. 1C, D). Altogether, the results indicate that mESCs tolerate GSK-3484862 for up to two weeks at concentrations up to 10 µM.

### Genes repressed by DNA methylation show derepression in GSK-3484862

We next assayed expression levels of seven germline genes (*Ctcfl, Magea4, Rhox1, Hormad1, Sohlh2, Dazl, Tuba3b*) and one transposon class (*Gln*) reported to be repressed by DNA methylation in mESCs[29-31]. As expected, all tested transcripts were highly upregulated in TKO cells (Fig. 2A, Supplementary Table 1). These transcripts were also upregulated after treatment with GSK-3484862, although often to a lesser extent than in TKO cells, potentially reflecting continued expression of DNMT3A and DNMT3B in the WT cells. GSK-3484862 produced by both companies showed similar levels of upregulation (Supplementary Table 1). Upregulation of target transcripts was observed in WT cells treated with either 2 µM or 10 µM GSK-3484862, starting as early as after two days of exposure (Figure 2), and all genes showed statistically significant upregulation at both concentrations in a one-sided t-test (Supplementary Table 2).

**Figure 2.**
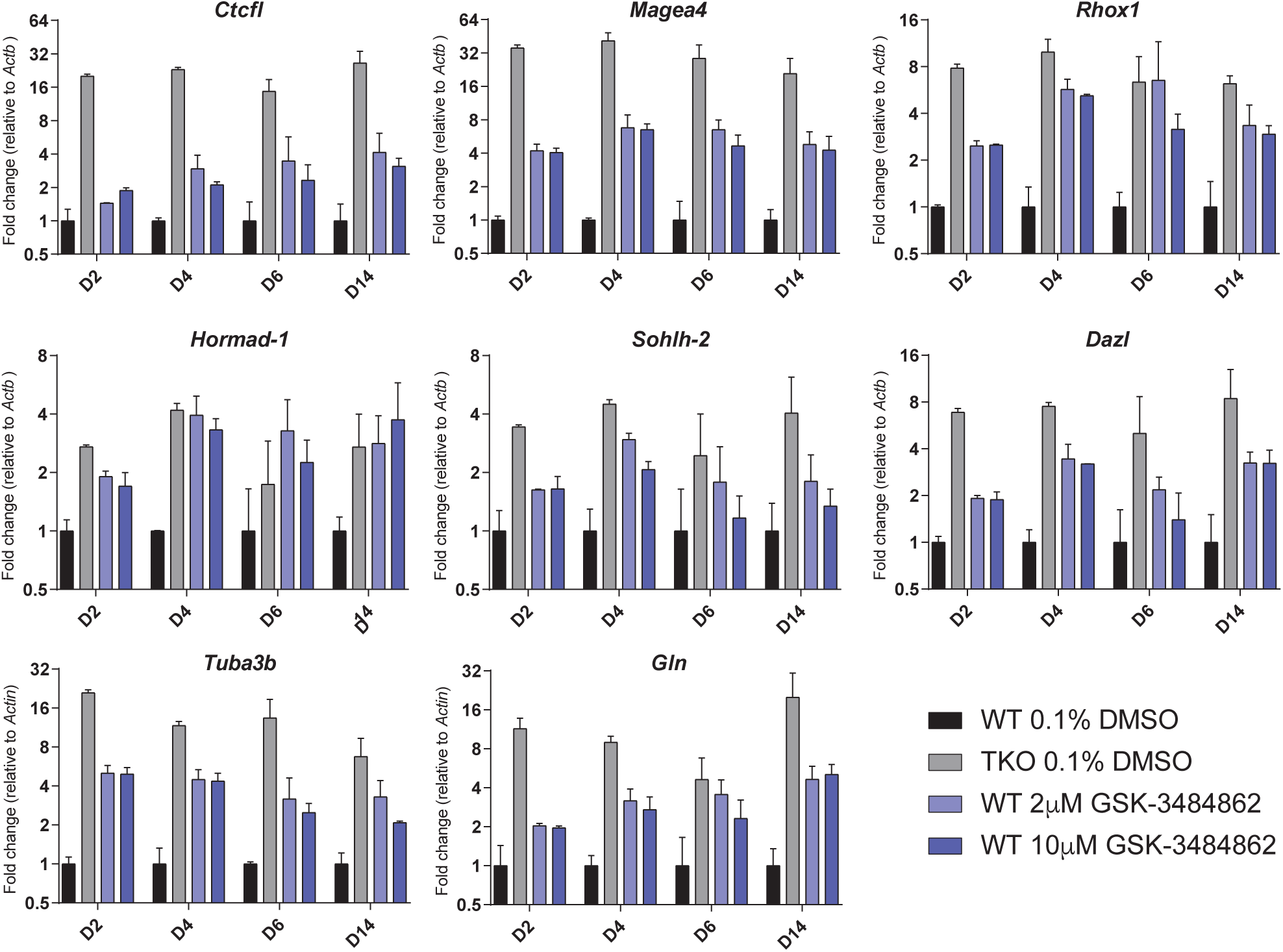
Expression of methylation-regulated transcripts in GSK-3484862 treated cells. RT-qPCR, normalised to Actb, of known methylation-regulated germline genes/transposons in cells treated with GSK-3484862 over the number of days indicated. N=2 – 6 biological replicates per sample. Mean and standard error are indicated.

### Severe toxicity from 5-azacytidine or decitabine treatment

WT and TKO cells were exposed to 5-azacytidine or decitabine for 48 hours. Even 0.1 µM concentrations of these compounds inflicted widespread cell death in WT cells (Fig 3A,B). TKO cells, which lack DNMT1 and thus will not form DNMT1-DNA adducts, appeared completely unaffected. After 1 – 2 days of recovery time, some WT ESC colonies of typical morphology were observed from the 0.1 µM and 0.3 µM 5-azacytidine and the 0.1 µM decitabine samples (Fig. 3A-D), and these were used for subsequent analysis.

**Figure 3.**
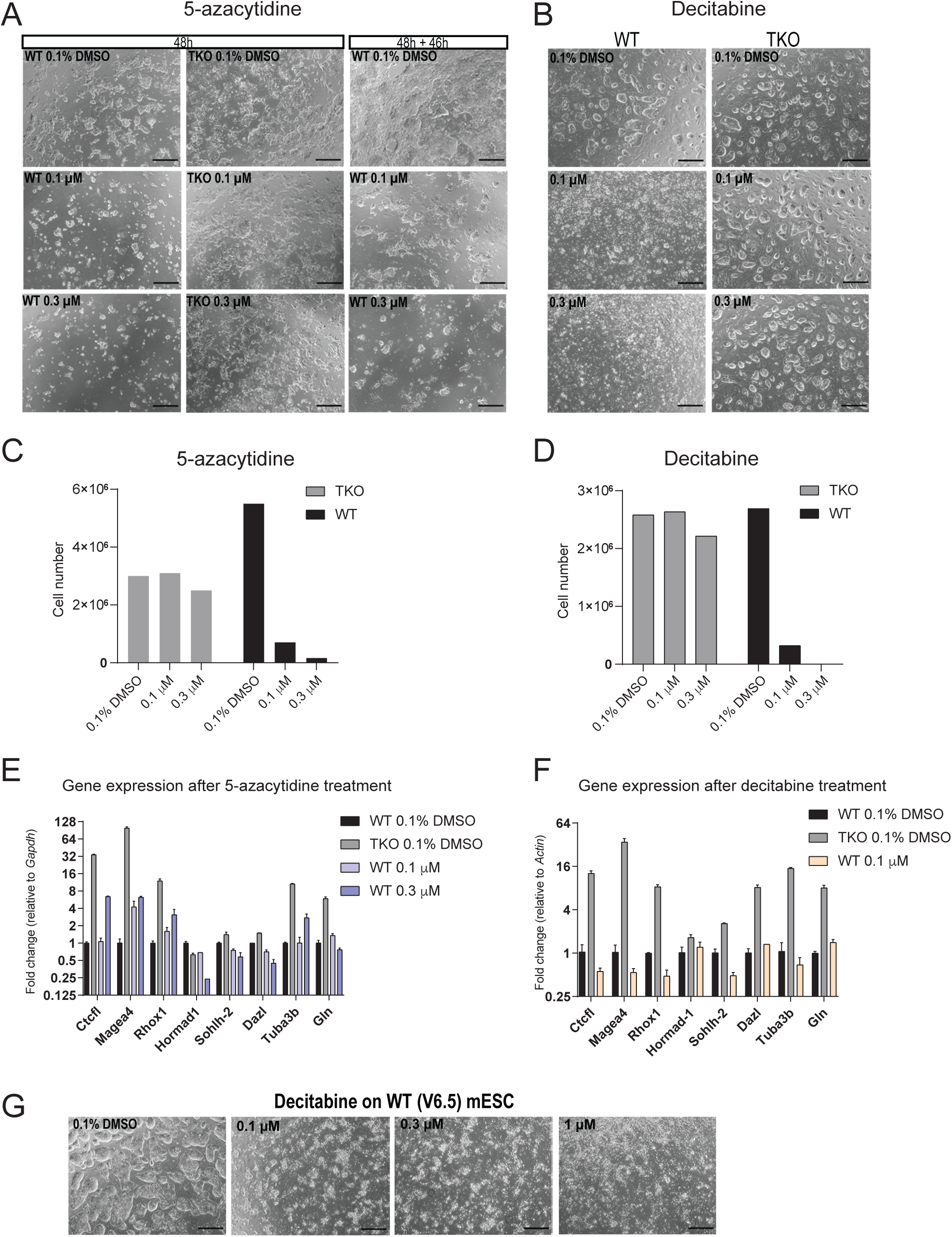
Treatment of mouse embryonic stem cells with 5-azanucleosides. **A** Brightfield images of WT and Dnmt1/3a/3b TKO cells after 48 hours of exposure to 0.1% DMSO, 0.1 or 0.3 µM 5-azacytidine (left columns) and WT cells 46 hours after the end of treatment with 5-azacytidine (right column). **B** Brightfield images taken of WT and TKO cells after 48 hours of exposure to 0.1% DMSO, 0.1 or 0.3 µM decitabine. To enhance survival, these cells were cultured on murine embryonic feeders (MEFs) rather than gelatin, hence the distinct morphology. **C, D** Cell numbers after treatment and recovery for 5-azacytidine **(C)** and decitabine **(D). E, F** Expression of methylated genes upon 5-azacytidine **(E)** or decitabine **(F)** treatment and recovery, normalized to Actb and Gapdh respectively. Mean and standard error of two technical replicates is indicated. **G** V6.5 mESCs after exposure to decitabine. These cells showed similar lethality to J1 WT mESCs.

WT cells treated with 0.3 µM 5-azacytidine showed upregulation of the genes *Tuba3b, Ctcfl, Magea4* and *Rhox1*, while there was no change in the expression of the other transcripts tested (Fig. 3E). Minimal or no upregulation of methylation-repressed transcripts was observed in WT cells treated with decitabine (Fig. 3F.)

Comparable decitabine-mediated toxicity was observed for WT mESCs from the genetic hybrid V6.5 line, confirming that this was not a cell line specific effect (Fig. 3G).

### Whole genome bisulfite sequencing showed dramatic demethylation in mESCs after GSK-3484862 treatment

Whole genome bisulfite sequencing (WGBS), even at low sequencing depth, provides an accurate quantitative estimate of total CpG methylation[32]. WT mESCs showed near 70% CpG methylation, dropping below 18% after six or 14 days of treatment with GSK-3484862, for both companies and at both concentrations (Fig. 4A). Strikingly, the methylation level of treated mESCs was almost identical to the published DNA methylation level of *Dnmt1* KO mESCs[29], consistent with total and specific inhibition of DNMT1. Mapping statistics and conversion efficiencies are shown in Supplementary Table 3.

**Figure 4.**
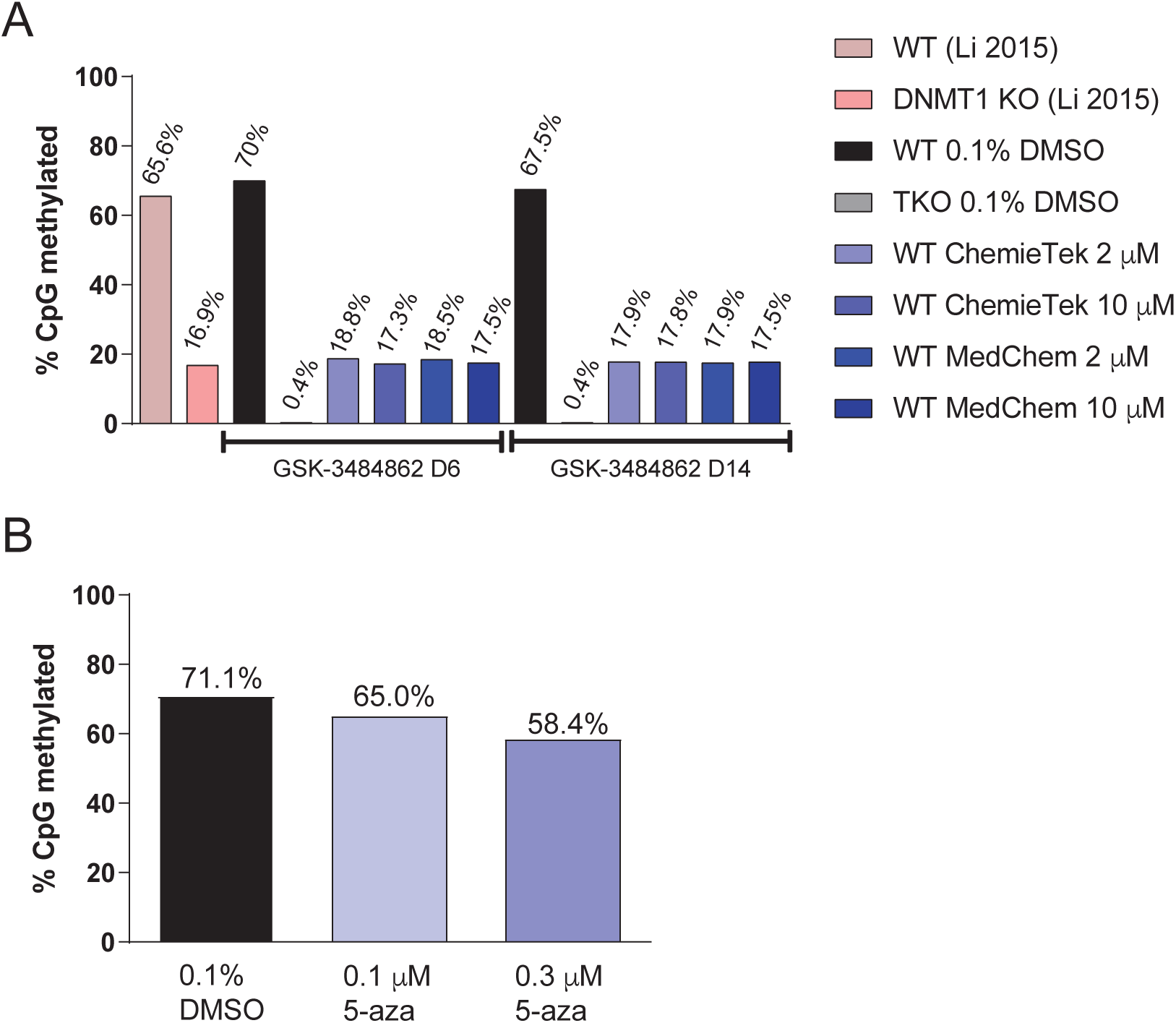
CpG methylation reduction after treatment with GSK-3484862 or 5-azacytidine. **A** Global CpG methylation levels of WT cells treated with 0.1% DMSO, Dnmt1/3a/3b TKO cells treated with 0.1% DMSO, and WT cells after 6 or 14 days of treatment with 2 µM or 10 µM GSK-3484862. Methylation levels of published Dnmt1 knockout cells are shown in comparison[29]. **B** Methylation levels in WT mESCs after treatment with 0.1% DMSO, 0.1 or 0.3 µM 5-azacytidine. WGBS = Whole-genome bisulfite sequencing.

Cells treated with 5-azacytidine showed far more modest reductions in 5-methylcytosine (Fig. 4B), but they may have regained some DNA methylation during the 46-hour recovery period after treatment.

## Discussion

From a research perspective, GSK-3484862 shows a great deal of promise. 5-azanucleosides have substantial non-specific toxicity and work within a narrow concentration band. We observed reactivation of some methylated genes after treatment with 0.3 µM 5-azacytidine but very few cells survived at this concentration, while 0.1 µM 5-azacytidine was inadequate to reactivate methylated genes. Furthermore, we had to allow two days for colonies of surviving cells to emerge in culture. By contrast, 2 µM or 10 µM GSK-3484862 reactivated methylated gene expression and produced only modest reduction in growth, most noticeable between days 6 and 10 of treatment. This reduced growth may reflect specific activity of the compound. When mESCs undergo demethylation mediated by treatment with MEK inhibitor, GSK3 inhibitor and high concentration of ascorbic acid, they have a burst of transposon expression and undergo reconfiguration of heterochromatin state approximately during this interval[31].

DNA methylation of the GSK-3484862-treated mESCs never fell below 17% regardless of the dosage used or time of treatment, and methylated genes were not reactivated to the extent observed in the TKO cells. This likely reflects the high level of DNMT3A and DNMT3B activity in mESCs, as evidenced by the similar level of DNA methylation in published *Dnmt1* KO mESCs[29]. The *Dnmt1* deficient or inhibited cells potentially reach an equilibrium in which methylation is constantly added by DNMT3A and DNMT3B and lost through replication and Tet-protein activity. Other cell types may respond to *Dnmt1* inhibition differently. Most somatic and cancer cells do not express such high levels of the *de novo* DNA methyltransferases and may not be able to maintain such high levels of DNA methylation in the absence of *Dnmt1* activity. At the same time, somatic or cancer cells may not survive dramatic DNA methylation loss[27, 33]. *Dnmt1* deficient and *Dnmt1/3a/3b* TKO mESCs are not viable upon differentiation, and DNMT1 becomes essential after implantation[6, 28]. This shift may reflect the fact that mESCs depend heavily on Trim28 to silence transposons, but upon differentiation of mESCs or uterine implantation of embryos, DNA methylation gains importance for transposon repression [7, 30, 34, 35]. Thus, researchers working with other cell types may well observe specific toxicity at lower doses. We also cannot rule out that non-specific toxicity may occur in some cell types, a result suggested by the GSK-3484862’s ability to halt mouse development at relatively low concentrations[25]. Nonetheless, this novel DNA methyltransferase inhibitor appears to be a substantial improvement over 5-azanucleosides and a promising research tool.

## Conclusions

GSK-3484862 mediates dramatic demethylation in murine embryonic stem cells. With regard to both activation of methylated genes and non-specific toxicity, GSK-3484862 performs far better than 5-azanucleosides.

## Methods

### Cell Culture

Mouse embryonic stem cells (mESC) lines used in this study included *Dnmt1/3a/3b* triple knockout [28] on a J1 background (129S4/SvJae), as well as wildtype J1 and V6.5 lines (C57BL/6 × 129S4/SvJae). Cells were cultured in Knockout-DMEM (Gibco) with 15% ES qualified FBS (Gibco), ESGRO mouse LIF 1000U/mL (Millipore), 1X PenStrep (Invitrogen), 1X Glutamax (Invitrogen), 55µM b-mercaptoethanol (Invitrogen), 1X non-essential amino acids (Invitrogen) and 1X Primocin (Invivogen). Cells were maintained at 37 °C and 5% CO_2_ on dishes coated with 0.1% gelatin (Millipore) and a layer of mitomycin C-treated mouse embryonic fibroblasts (MEFs) (Thermofisher Scientific).

### Determining GSK-3484862 toxicity

An assay to determine the optimal concentration and toxicity of GSK-3484862 (ChemieTek) was performed using J1-line WT and DNMT TKO mESCs. 30,000 cells were seeded in 24-well plates pre-coated with 0.1% gelatin. The next day, medium was changed to fresh mESC medium or medium containing DMSO (0.1% or 1%) for the following concentrations of GSK-3484862: 2 pM, 20 pM, 200 pM, 2 nM, 20 nM, 200 nM, 2 µM, 20 µM (in 0.1% DMSO) and 200 µM (in 1% DMSO). The medium was refreshed every day for the next six days, after which cell morphology was assessed, followed by cell dissociation with 0.05% Trypsin-EDTA (Gibco) for cell counting.

Next, the demethylation efficacy and long-term cytotoxicity of GSK-3484862 from two companies, ChemieTek and MedChemExpress, was evaluated in duplicate experiments. To improve solubility, after resuspension in DMSO, GSK-3484862 was subjected to ultrasonication. GSK-3484862 from both companies were sonicated for six minutes at 42 kHz in an ultrasonic water bath (Sper Scientific). Still, drug precipitation was observed for concentrations at or above 20 µM in media, and therefore an upper concentration of 10 µM was chosen. WT and DNMT TKO cells were seeded in 12-well plates pre-coated with 0.1% gelatin and had 0.1% DMSO, 2 µM or 10 µM GSK3484862 added in medium + LIF from day zero. The medium was refreshed every day and cells were counted using the Countess II FL instrument and passaged every 2-3 days for the next 14 days.

### 5-azacytidine assay

We found that WT mESCs died if treated with 5-azacytidine immediately after plating (data not shown). Therefore, we plated cells at low density (1.4 × 10^4^ cells/ cm^2^) and added 0.1% DMSO, 0.1 or 0.3 µM of the drug 24 hours later. WT and DNMT TKO cells were exposed to 5-azacytidine for 48h, with a media change at the 24-hour timepoint to replenish 5-azacytidine. The cultures were then maintained in media without the drug for 46 hours. Cells were then dissociated with 0.05% Trypsin-EDTA, counted, and collected for subsequent analysis. Because of their higher survival and consequent higher density, DNMT TKO cells were harvested after only 24 hours recovery time.

### Decitabine assay

Concentrations of decitabine as low as 0.1 µM proved lethal to WT mESCs plated on gelatin (data not shown), so mESCs were seeded at 1.4 × 10^4^ cells/ cm^2^ on a monolayer of MEF feeder cells to enhance survival. 24 hours later, 0.1% DMSO or decitabine at concentrations of 0.1 µM, 0.3 µM or 1 µM was added and cells were treated for 48 hours, with a media change at the 24-hour timepoint. Cells were allowed to recover for another 24 hours and then harvested. Because of the low viability of the WT J1 line mESCs, the experiment was repeated with V6.5 mESCs, a robust line on a hybrid genetic background (C57BL/6 × 129S4/SvJae).

### Cell imaging and counting

All cells in this paper were imaged with an EVOS M5000 microscope (Invitrogen) and counted using Countess II FL Automated Cell Counter (Invitrogen). Images were processed using Adobe Photoshop and ImageJ, and graphs were created using GraphPad Prism software.

### DNA and RNA extraction

Cell pellets were snap frozen and kept at –80 °C until extraction. Samples from the GSK-3484862 and 5-azacytidine treated assays had RNA and DNA isolated simultaneously using the AllPrep DNA/RNA kit (Qiagen), whereas only RNA was isolated from decitabine samples by using the RNAzol total RNA protocol (Sigma). DNA and RNA concentrations were measured with the Qubit dsDNA and RNA HS Assay Kits respectively (ThermoFisher).

### Quantitative RT-PCR

500 ng of RNA was used for cDNA synthesis using Froggabio SensiFAST cDNA synthesis kit and following the manufacturer’s instructions. The qPCR reaction was performed using PowerUp SYBR green mix (Invitrogen) containing cDNA generated from the equivalent of 5ng RNA and 0.5 μM of primer mix (forward and reverse) in a final reaction of 6 µL per duplicate. Quantification and analysis were performed with the QuantStudio5 instrument (Applied Biosystems), and the cycling conditions were: (50°C 2 min, 95°C 20 s, 55x (95°C 3 s, 60°C 30 s), 95°C 1 s). The following primers were used:

*Actin*: F:5’-ACTGGGACGACATGGAGAAG -3’ R:5’-GGGGTGTTGAAGGTCTCAAA -3’

*Gapdh*: F:5’-CATCAAGAAGGTGGTGAAGC -3’ R:5’-GGGAGTTGCTGTTGTAAGTCG -3’

*Tuba3b*: F:5’-AGGAAGATGCAGCCAACAATTA -3’ R:5’-TGCACAGATCGGCCAGTTT -3’

*Ctcfl*: F:5’-GCCTTCAGCATTGCGTGAC -3’ R:5’-AGCAGGTGAAAATGTATCCGC -3’

*Magea4*: F:5’-GTCTCTGGCATTGGCATGATAG -3’ R:5’-GCTTACTCTGAACATCAGTCAGC -3’

*Rhox1*: F:5’-CCGGTTTTCTGGAGTATGAGAGA -3’ R:5’-CCAGCCGTTTTCTGTCTTGTG -3’

*Hormad*-1: F:5’-TGAAAACTCTGGAGCTTCTGAAA -3’ R:5’-ACTGACTAACTGTTCAACCTGACA -3’

*Sohlh*-2: F:5’-CCATCGAGCTGTTCCTTCCA -3’ R:5’-GGAATACACGTTCAGGCCCC -3’

*Dazl*: F:5’-GTGGCTTCTGCTCCACCTTCG -3’ R:5’-CCTTGACTTGTGGTTGCTGA -3’

*Gln*: F:5’-CGTAAGGACCCTAGTGGCTG -3’ R:5’-GCACTCACTCTTCTTCACTCTG -3’

The *Dazl* and *Gln* primers were taken from published sources[30].

### Analysis of qRT-PCR data

To calculate which genes showed statistically significant upregulation relative to control cells, expression of each gene in the DNMT TKO or GSK-3484862 cells was normalized to the control or control lines in the same experimental replicate and time-course. Data from cells treated with GSK-3484862 from the two manufacturers (ChemieTek and MedChemExpress) was combined. A paired, one tailed t-test was conducted, pairing control cells with treated cells for each time point and calculating one p-value for each GSK-3484862 concentration across all timepoints.

### Whole-genome bisulfite library (WGBS) preparation

Genomic DNA (500 ng) with spiked-in lambda DNA (1.25 ng) (NEB) was sheared using an M220 ultrasound sonicator (Covaris) to an average size of 350 bp. The size of the DNA fragments was confirmed by electrophoresis on an 1.5% agarose gel. 200 ng of sheared DNA was subjected to bisulfite conversion using EZ DNA Methylation-Gold Kit (Zymo Research) as per manufacturer’s instructions. Libraries were prepared using 50-100 ng of bisulfite-converted DNA and the Accel-NGS Methyl-Seq DNA Library prep kit (Swift Biosciences) according to manufacturer’s instructions, with 8-11 cycles of PCR-amplification. The integrity of the libraries was assessed using agarose gel electrophoresis.

Libraries were sequenced on an Illumina HiSeq 4000 instrument at the Michael Smith Genome Sciences Centre at BC Cancer.

### WGBS analysis

Raw reads were quality checked with FastQC and adapters at either end were trimmed with the Trim Galore! (v0.6.7) software. Reads were then aligned to the mm10 reference genome using Bismark (v0.21.0). In cases where spike-in DNA was used to quality check bisulfite conversion rates, reads were aligned either to pUC19 or lambda DNA, as specified (Supplementary Table 3). PCR duplicates were removed using the deduplicate command in Bismark. DNA methylation was subsequently calculated using the methylation extractor command in Bismark. In all cases, default parameters were used.

## Acknowledgements

We thank Canada’s Michael Smith Genome Sciences Centre at BC Cancer for dedicated service. We thank Masaki Okano (Kumamoto University) and Amander Clark lab (UCLA) for providing embryonic stem cell lines. We thank Colleen Russett (McGill University) for preliminary work on this project. We thank Daniel Sapozhnikov (McGill University), Chuck Haggerty and Alexander Meissner (Max Planck Institute), and Melissa Pappalardi (GlaxoSmithKline) for helpful information.

## Contributions

NAP, DS and JS conducted experiments. IH analyzed bisulfite sequencing data. NAP and WP wrote the manuscript. CM and WP supervised research.

## Figure Captions

**Supplementary Table 1. qRT-PCR data for all samples**.

**Supplementary Table 2. P-values for upregulation of indicated genes**.

**Supplementary Table 3. Mapping and conversion statistics for Whole Genome Bisulfite Sequencing data used in this publication**.

## References

1. Schubeler D: Function and information content of DNA methylation. Nature 2015, 517(7534):321–326.

2. Baubec T, Ivanek R, Lienert F, Schubeler D: Methylation-dependent and -independent genomic targeting principles of the MBD protein family. Cell 2013, 153(2):480–492.

3. Okano M, Bell DW, Haber DA, Li E: DNA methyltransferases Dnmt3a and Dnmt3b are essential for de novo methylation and mammalian development. Cell 1999, 99(3):247–257.

4. Goyal R, Reinhardt R, Jeltsch A: Accuracy of DNA methylation pattern preservation by the Dnmt1 methyltransferase. Nucleic Acids Res 2006, 34(4):1182–1188.

5. Wang Q, Yu G, Ming X, Xia W, Xu X, Zhang Y, Zhang W, Li Y, Huang C, Xie H et al: Imprecise DNMT1 activity coupled with neighbor-guided correction enables robust yet flexible epigenetic inheritance. Nat Genet 2020, 52(8):828–839.

6. Li E, Bestor TH, Jaenisch R: Targeted mutation of the DNA methyltransferase gene results in embryonic lethality. Cell 1992, 69(6):915–926.

7. Dahlet T, Argueso Lleida A, Al Adhami H, Dumas M, Bender A, Ngondo RP, Tanguy M, Vallet J, Auclair G, Bardet AF et al: Genome-wide analysis in the mouse embryo reveals the importance of DNA methylation for transcription integrity. Nat Commun 2020, 11(1):3153.

8. Smith ZD, Shi J, Gu H, Donaghey J, Clement K, Cacchiarelli D, Gnirke A, Michor F, Meissner A: Epigenetic restriction of extraembryonic lineages mirrors the somatic transition to cancer. Nature 2017, 549(7673):543–547.

9. Da Costa EM, McInnes G, Beaudry A, Raynal NJ: DNA Methylation-Targeted Drugs. Cancer J 2017, 23(5):270–276.

10. Sorm F, Piskala A, Cihak A, Vesely J: 5-Azacytidine, a new, highly effective cancerostatic. Experientia 1964, 20(4):202–203.

11. Sorm F, Vesely J: Effect of 5-aza-2’-deoxycytidine against leukemic and hemopoietic tissues in AKR mice. Neoplasma 1968, 15(4):339–343.

12. Jones PA, Taylor SM: Cellular Differentiation, Cytidine Analogs, and DNA Methylation. Cell 1980, 20:85–93.

13. Stresemann C, Lyko F: Modes of action of the DNA methyltransferase inhibitors azacytidine and decitabine. Int J Cancer 2008, 123(1):8–13.

14. Schermelleh L, Spada F, Easwaran HP, Zolghadr K, Margot JB, Cardoso MC, Leonhardt H: Trapped in action: direct visualization of DNA methyltransferase activity in living cells. Nat Methods 2005, 2(10):751–756.

15. Patel K, Dickson J, Din S, Macleod K, Jodrell D, Ramsahoye B: Targeting of 5-aza-2’-deoxycytidine residues by chromatin-associated DNMT1 induces proteasomal degradation of the free enzyme. Nucleic Acids Res 2010, 38(13):4313–4324.

16. Jabbour E, Issa JP, Garcia-Manero G, Kantarjian H: Evolution of decitabine development: accomplishments, ongoing investigations, and future strategies. Cancer 2008, 112(11):2341–2351.

17. Li LH, Olin EJ, Buskirk HH, Reineke LM: Cytotoxicity and mode of action of 5-azacytidine on L1210 leukemia. Cancer Res 1970, 30(11):2760–2769.

18. Lee TT, Karon MR: Inhibition of protein synthesis in 5-azacytidine-treated HeLa cells. Biochem Pharmacol 1976, 25(15):1737–1742.

19. Schaefer M, Hagemann S, Hanna K, Lyko F: Azacytidine inhibits RNA methylation at DNMT2 target sites in human cancer cell lines. Cancer Res 2009, 69(20):8127–8132.

20. Farlik M, Sheffield NC, Nuzzo A, Datlinger P, Schonegger A, Klughammer J, Bock C: Single-cell DNA methylome sequencing and bioinformatic inference of epigenomic cell-state dynamics. Cell Rep 2015, 10(8):1386–1397.

21. Maurano MT, Wang H, John S, Shafer A, Canfield T, Lee K, Stamatoyannopoulos JA: Role of DNA Methylation in Modulating Transcription Factor Occupancy. Cell Rep 2015, 12(7):1184–1195.

22. Hackett JA, Reddington JP, Nestor CE, Dunican DS, Branco MR, Reichmann J, Reik W, Surani MA, Adams IR, Meehan RR: Promoter DNA methylation couples genome-defence mechanisms to epigenetic reprogramming in the mouse germline. Development 2012, 139(19):3623–3632.

23. Adams ND, Benowitz AB, Benede MLR, Evans KA, Fosbenner DT, King BW, Li M, Luengo JI, Miller WH, Reif AJ et al:. Substituted Pyridines as Inhibitors of DNMT1. In., vol. WO/2017/216727; 2017.

24. Gilmartin AG, Groy A, Gore ER, Atkins C, Long ER, Montoute MN, Wu Z, Halsey W, McNulty DE, Ennulat D et al: In vitro and in vivo induction of fetal hemoglobin with a reversible and selective DNMT1 inhibitor. Haematologica 2021, 106(7):1979–1987.

25. Haggerty C, Kretzmer H, Riemenschneider C, Kumar AS, Mattei AL, Bailly N, Gottfreund J, Giesselmann P, Weigert R, Brandl B et al: Dnmt1 has de novo activity targeted to transposable elements. Nat Struct Mol Biol 2021, 28(7):594–603.

26. Ficz G, Hore TA, Santos F, Lee HJ, Dean W, Arand J, Krueger F, Oxley D, Paul YL, Walter J et al: FGF signaling inhibition in ESCs drives rapid genome-wide demethylation to the epigenetic ground state of pluripotency. Cell Stem Cell 2013, 13(3):351–359.

27. Jackson-Grusby L, Beard C, Possemato R, Tudor M, Fambrough D, Csankovszki G, Dausman J, Lee P, Wilson C, Lander E et al: Loss of genomic methylation causes p53-dependent apoptosis and epigenetic deregulation. Nat Genet 2001, 27(1):31–39.

28. Tsumura A, Hayakawa T, Kumaki Y, Takebayashi S, Sakaue M, Matsuoka C, Shimotohno K, Ishikawa F, Li E, Ueda HR et al: Maintenance of self-renewal ability of mouse embryonic stem cells in the absence of DNA methyltransferases Dnmt1, Dnmt3a and Dnmt3b. Genes Cells 2006, 11(7):805–814.

29. Li Z, Dai H, Martos SN, Xu B, Gao Y, Li T, Zhu G, Schones DE, Wang Z: Distinct roles of DNMT1-dependent and DNMT1-independent methylation patterns in the genome of mouse embryonic stem cells. Genome Biol 2015, 16:115.

30. Karimi MM, Goyal P, Maksakova IA, Bilenky M, Leung D, Tang JX, Shinkai Y, Mager DL, Jones S, Hirst M et al: DNA methylation and SETDB1/H3K9me3 regulate predominantly distinct sets of genes, retroelements, and chimeric transcripts in mESCs. Cell Stem Cell 2011, 8(6):676–687.

31. Walter M, Teissandier A, Perez-Palacios R, Bourc’his D: An epigenetic switch ensures transposon repression upon dynamic loss of DNA methylation in embryonic stem cells. Elife 2016, 5.

32. Bewick AJ, Hofmeister BT, Lee K, Zhang X, Hall DW, Schmitz RJ: FASTmC: A Suite of Predictive Models for Nonreference-Based Estimations of DNA Methylation. G3 (Bethesda) 2015, 6(2):447–452.

33. Chen T, Hevi S, Gay F, Tsujimoto N, He T, Zhang B, Ueda Y, Li E: Complete inactivation of DNMT1 leads to mitotic catastrophe in human cancer cells. Nat Genet 2007, 39(3):391–396.

34. Rowe HM, Jakobsson J, Mesnard D, Rougemont J, Reynard S, Aktas T, Maillard PV, Layard-Liesching H, Verp S, Marquis J et al: KAP1 controls endogenous retroviruses in embryonic stem cells. Nature 2010, 463(7278):237–240.

35. Hutnick LK, Huang X, Loo TC, Ma Z, Fan G: Repression of retrotransposal elements in mouse embryonic stem cells is primarily mediated by a DNA methylation-independent mechanism. J Biol Chem 2010, 285(27):21082–21091.

